# Compared with SARS-CoV2 wild type’s spike protein, the SARS-CoV2 omicron’s receptor binding motif has adopted a more SARS-CoV1 and/or bat/civet-like structure

**DOI:** 10.1101/2021.12.14.472585

**Authors:** Michael O. Glocker, Kwabena F.M. Opuni, Hans-Juergen Thiesen

## Abstract

Our study focuses on free energy calculations of SARS-CoV2 spike protein receptor binding motives (RBMs) from wild type and variants-of-concern with particular emphasis on currently emerging SARS- CoV2 omicron variants of concern (VOC). Our computational free energy analysis underlines the occurrence of positive selection processes that specify omicron host adaption and bring changes on the molecular level into context with clinically relevant observations. Our free energy calculations studies regarding the interaction of omicron’s RBM with human ACE2 shows weaker binding to ACE2 than alpha’s, delta’s, or wild type’s RBM. Thus, less virus is predicted to be generated in time per infected cell. Our mutant analyses predict with focus on omicron variants a reduced spike-protein binding to ACE2-receptor protein possibly enhancing viral fitness / transmissibility and resulting in a delayed induction of danger signals as trade-off. Finally, more virus is produced but less per cell accompanied with delayed Covid-19 immunogenicity and pathogenicity. Regarding the latter, more virus is assumed to be required to initiate inflammatory immune responses.

## Amino acid sequence alignments point to a shift in RBM characteristics

Within the receptor binding domain (RBD; aa319 to aa541) of the wild type (wt) SARS-CoV2 spike protein, the amino acid sequence stretch aa437 to aa508 encompasses the receptor binding motif (RBM) [1]. Amino acid residue exchanges have been observed at distinct RBM positions with all variants of concern (VOCs) [2]. The newly reported omicron VOC carries ten exchanged amino acids (Supplementary Figure 1) in its RBM of which five (K440, S446, N447, K478, and A484) are also found in SARS-CoV1-, in bat-, and/or in civet-derived RBMs at the respective positions and through which omicron’s RBM can be distinguished from that of SARS-CoV2’s wt [3]. 3D structures of SARS-CoV1 [4] and SARS-COV2 wt [1] are from X-ray data, whereas the RBM structure of SARS-COV2 o has been modeled by alphafold [5]. From the remaining five exchanged amino acid residues, four (K493, S496, R498, and H505) are unique to omicron which further differentiates omicron’s from wt’s SARS-CoV2 spike protein when comparing the here assembled seven RBMs (Table 1). Importantly, omicron encodes for Y501 which strengthened binding already in alpha, beta, and gamma VOCs [6]. Of note, bat RBMs (BM48-31 and Rp3) do not bind to human ACE2 [7] and SARS-CoV1 binds to human ACE2 with lower affinity than SARS-CoV2 wt [8].

**Table 1:**
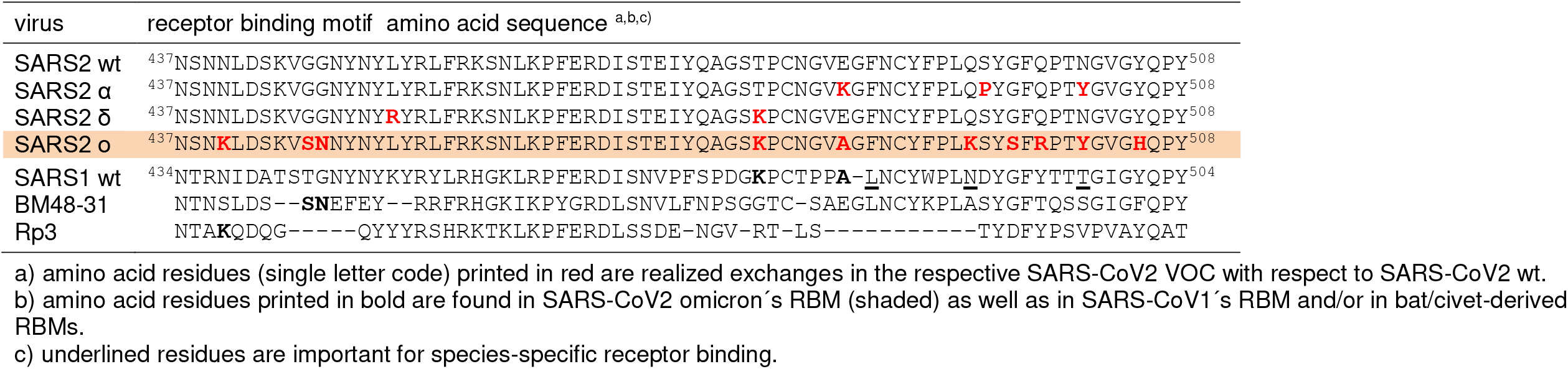
Amino acid sequence alignments of corona virus spike protein receptor binding motifs.

In detail: residue K478 has been designated the decisive amino acid exchange in delta’s RBM [2]. Omicron has kept K478 which, like residues K440, S446, and N447 (all three are rarely seen in other variants [9]), lend omicron “non-SARS-CoV2” characteristics; *e*.*g*. K478 matches with K465 in the RBM of SARS-CoV1. With expressing A484 omicron avoided the in alpha and in other VOCs found receptor binding-weakening E484 residue [10]. The A484 matching residue from SARS-CoV1’s RBM is A471 which is located adjacent to L472, one of the amino acid residues, which is in direct contact with human ACE2 and which has been assigned as important for species specific binding [4]. Residue K493 in omicron’s RBM is positioned where N479 is found in SARS-CoV1’s RBM. N479 makes direct contact with human ACE2 and is considered to be responsible for species-specific binding as well. An N479K exchange resulted in steric hindrance and in weakening of RBD-binding to human ACE2 [11]. S496 and R498 are rare RBM mutations and adverse effects on binding can be estimated for R498 as opposed to Q498 on wt’s RBM because of charge repulsion [4]. Y501 was considered to strengthen binding to human ACE2 considerably with respect to SARS-CoV2 wt and was assumed to also increase virus replication rates [12-14]. At last, H505 from SARS-CoV2 omicron is located where Y491 is placed in SARS-CoV1. H505 replaces Y505 of SARS-CoV2 wt’s RBD and of other VOCs, respectively [3]. Y505 is directly involved in binding to human ACE2 and from the physico-chemical properties of histidine vs. tyrosine one can conclude that binding of omicron’s RBD to ACE2 would not be positively affected by this exchange.

In sum, because of the multitude of amino acid exchanges, the interaction of omicron’s RBM with human ACE2 is assumed to be weaker than that of alpha’s, delta’s, or wt’s RBM with human ACE2.

## Free energy calculations indicate weaker receptor binding of omicron’s RBM

To substantiate our hypothesis of weaker interactions between omicron’s RBM and ACE2 as compared to wt, we performed free energy difference calculations [15] for the SARS-CoV2-derived RBMs (wt vs. alpha or delta or omicron) when bound to human ACE2 (Table 2). We then compared these binding differences to respective RBM - DPP-IV interactions [16]. Since human DPP-IV is considered not to function as a receptor for SARS-CoV2 *in vivo* [7], calculations of free energy differences of RBM - DPP-IV complexes served as controls. Of note, the MERS virus uses DPP-IV as a receptor and SARS-CoV2 wt has been assumed to as well being able to bind to DPP-IV [17].

**Table 2:**
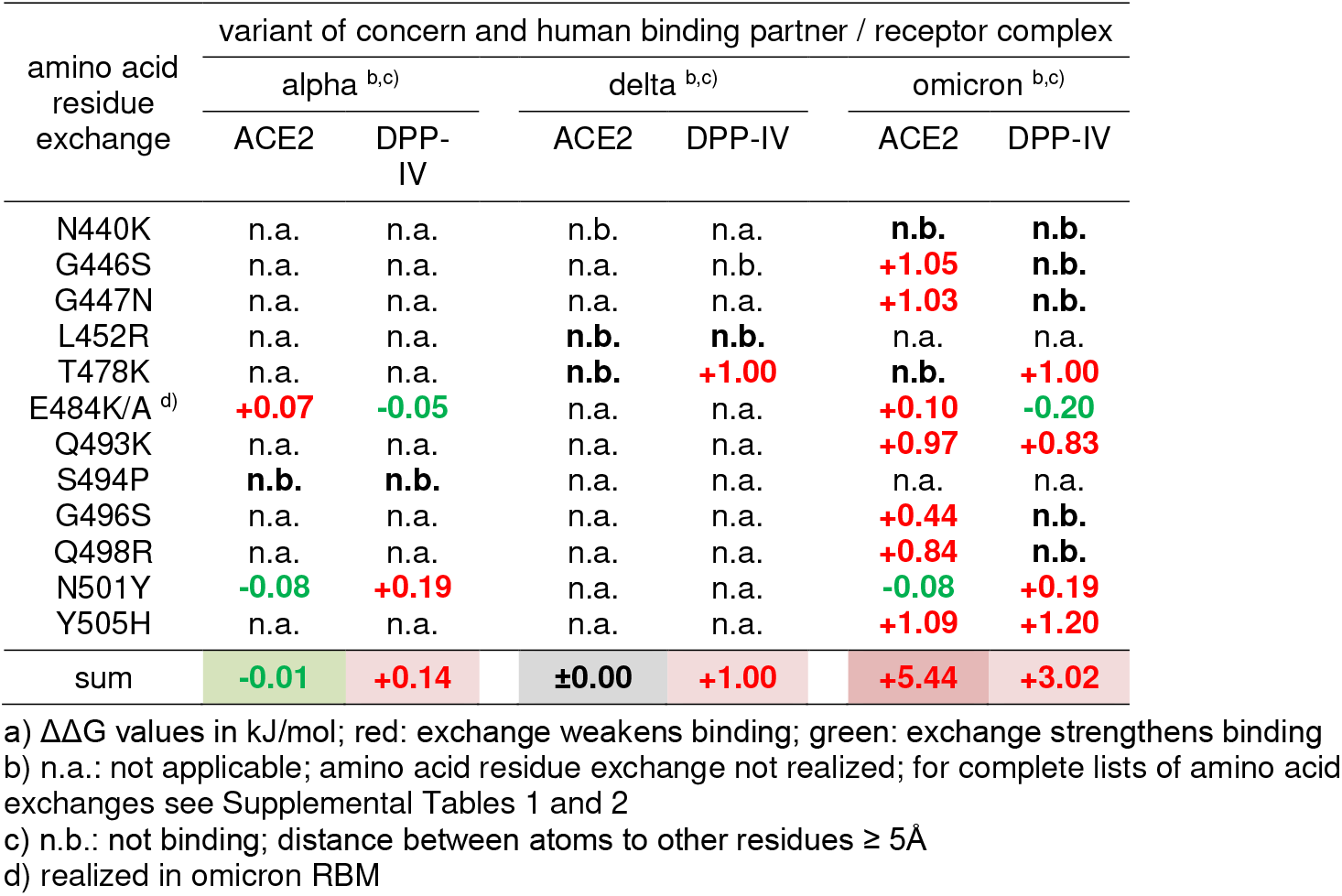
Spike protein receptor binding motif amino acid exchanges and changes of free energies with human ACE2 binding or human DPP-IV binding ^a)^.

| amino acid<br>residue<br>exchange | variant of concern and human binding partner / receptor complex |  |  |  |  |  |
| --- | --- | --- | --- | --- | --- | --- |
|  | alpha <sup>b,c)</sup> |  | delta <sup>b,c)</sup> |  | omicron <sup>b,c)</sup> |  |
|  | ACE2 | DPP-IV | ACE2 | DPP-IV | ACE2 | DPP-IV |
| N440K | n.a. | n.a. | n.b. | n.a. | **n.b.** | **n.b.** |
| G446S | n.a. | n.a. | n.a. | n.b. | **+1.05** | **n.b.** |
| G447N | n.a. | n.a. | n.a. | n.a. | **+1.03** | **n.b.** |
| L452R | n.a. | n.a. | **n.b.** | **n.b.** | n.a. | n.a. |
| T478K | n.a. | n.a. | **n.b.** | **+1.00** | **n.b.** | **+1.00** |
| E484K/A <sup>d)</sup> | **+0.07** | **-0.05** | n.a. | n.a. | **+0.10** | **-0.20** |
| Q493K | n.a. | n.a. | n.a. | n.a. | **+0.97** | **+0.83** |
| S494P | **n.b.** | **n.b.** | n.a. | n.a. | n.a. | n.a. |
| G496S | n.a. | n.a. | n.a. | n.a. | **+0.44** | **n.b.** |
| Q498R | n.a. | n.a. | n.a. | n.a. | **+0.84** | **n.b.** |
| N501Y | **-0.08** | **+0.19** | n.a. | n.a. | **-0.08** | **+0.19** |
| Y505H | n.a. | n.a. | n.a. | n.a. | **+1.09** | **+1.20** |
| sum | **-0.01** | **+0.14** | **±0.00** | **+1.00** | **+5.44** | **+3.02** |

a) $\Delta\Delta G$ values in kJ/mol; red: exchange weakens binding; green: exchange strengthens binding
b) n.a.: not applicable; amino acid residue exchange not realized; for complete lists of amino acid exchanges see Supplemental Tables 1 and 2
c) n.b.: not binding; distance between atoms to other residues $\geq 5\text{\AA}$
d) realized in omicron RBM

According to ΔΔG calculations on the respective amino acid exchanges and their contributions to receptor binding, one observes that with respect to the wt RBM the alpha RBM is the only one that achieved slightly stronger binding to human ACE2 when summing up all amino acid residue energy differences which arise from the respective single amino acid exchanges. The delta RBM neither gained nor lost binding strength compared to wt RBM – ACE2 binding. Surprisingly, the omicron RBM – ACE2 complex is energetically less favored (ΔΔG: +5.44 kJ/mol) than the complex between wt RBM and human ACE2 which means that the omicron’s RBM binding to ACE2 is weakened with respect to ACE2 binding of either the wt or the alpha or the delta RBM. Noteworthy, the presence of N417, though located outside omicron’s RBM, is known to reduce ACE2 binding [12,17] which correlates with our calculations where N417 affords an increase of free binding energy as compared to K417 of SARS-CoV2 wt’s RBM (ΔΔG +0.49 kJ/mol) as well as to DPP-IV (ΔΔG +0.64 kJ/mol); see Supplementary Table 1.

By contrast, ΔΔG value differences of RBD – DPP-IV binding of all VOCs showed that all their respective complexes were bound with weaker forces than that of wt. as well as to DPP-IV (Table 2). Of note, N417K exchange weakens binding to DPP-IV even more (ΔΔG +0.64 kJ/mol); see Supplementary Table 2. This stands in agreement with the observation that SARS-CoV2 wt uses human ACE2 as entry into host cells rather than DPP-IV [4]. Interestingly, omicron RBM binding with human DPP-IV requests a smaller increase in free energy (ΔΔG: +3.66 kJ/mol) than that of omicron RBM binding with human ACE2 (ΔΔG: +5.93 kJ/mol) when taking the N417k exchange into account. It remains to be investigated whether such a free energy difference was large enough to cause omicron to switch host receptors, hence, to possibly alter tropism and to eventually afford different disease symptoms.

## A molecular perspective on transmissibility and mutual disease outcome

The weaker binding of the spike protein to its receptor will slow down virus uptake into cells and, hence, elicit fewer danger signals, thereby retarding innate immune response [18] which over time might result in higher viral load in the upper respiratory tract. Also of interest, outside its RBM the omicron spike protein carries the N679K, P681H, N679K, D614G exchanges [19,20] which may assist in enhancing transmissibility [21,22].

On the other hand, Covid-19 is considered a result of an overacting immune response mostly affecting the lower respiratory tract [23]. It is tempting to speculate whether omicron’s assumed weaker RBM – ACE2 binding with respect to those of wt or alpha or delta contributed as well to clinical observations of less severe disease outcome upon SARS-CoV2 omicron infection as compared to infections with other SARS-CoV2 VOCs. Hitherto reported disease severity upon infections with SARS-CoV2 omicron, though yet anecdotal, has been graded mild or asymptomatic [24].

From the here outlined molecular perspectives, it seems plausible that SARS-CoV2 omicron fulfilled some key criteria of a host-adapted virus variant with high contagion potential and perhaps less severe disease outcome [25]. Particularly after monovalent vaccine administration, SARS-CoV2 omicron might challenge a human’s post-immunized waning antibody / B-cell responses to induce a more general and long lasting immunity by extending protective antibody repertoires and by simultaneously enhancing T-cell mediated immunity, thereby ultimately preparing an individual to defeat more pathogenic virus variants in the future.

## Funding

The authors declare that no funding sources have been engaged for writing this manuscript.

## Acknowledgements

The authors declare that no external help has been engaged for writing this manuscript.

## Conflict of Interest

The authors declare no conflicts of interest.

---

**Supplemental Figure 1:**
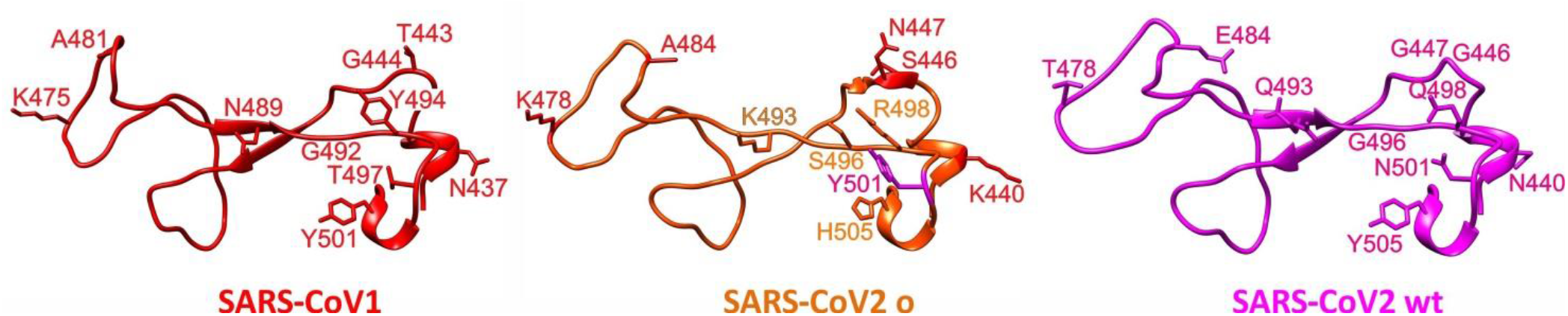
Structure comparisons of SARS-CoV receptor binding motifs. 3D structures (cartoon views) of SARS-CoV1 [4] and SARS-COV2 wt [1] are from X-ray data, whereas the RBM structure of SARS-COV2 o has been modeled by alphafold [5,6]. Mutated amino acid residues from SARS-COV2 o and their respective counterparts in SARS-CoV1 or SARS-CoV2 wt (labeled) are shown as stick models.

**Supplemental Table 1:** Effects of SARS-CoV2 RBD-exchanged amino acid residues on strengths of interaction with ACE2.

| amino acid<br>residues in<br>wt RBD <sup>b)</sup> | $\Delta\Delta G$ values in kcal / mol <sup>a)</sup> | | | | | | | | | | | | | | | | | | | |
| --- | --- | --- | --- | --- | --- | --- | --- | --- | --- | --- | --- | --- | --- | --- | --- | --- | --- | --- | --- | --- |
|  | A | C | D | E | F | G | H | I | K | L | M | N | P | Q | R | S | T | V | W | Y |
| K417 | +1.01 | +0.83 | +0.38 | +0.51 | +0.11 | +0.74 | +0.05 | +0.28 | $\pm 0.00$ | -0.00 | -0.23 | +0.49 | +0.18 | +0.47 | +0.23 | +0.87 | +0.51 | +0.79 | -0.24 | -0.10 |
| G446 | +1.27 | +0.93 | +1.05 | +1.44 | +1.17 | $\pm 0.00$ | +0.89 | +1.55 | +1.22 | +1.37 | +1.33 | +0.79 | +1.49 | +1.27 | +1.13 | +1.05 | +1.32 | +1.32 | +1.27 | +1.11 |
| G447 | +0.94 | +0.92 | +1.20 | +1.40 | +0.96 | $\pm 0.00$ | +0.84 | +1.01 | +1.22 | +1.09 | +0.88 | +1.03 | +1.15 | +1.09 | +1.08 | +1.01 | +1.04 | +1.08 | +0.79 | +0.97 |
| Y449 | +0.98 | +1.04 | +0.74 | +0.95 | +0.36 | +0.89 | +0.46 | +0.63 | +0.76 | +0.59 | +0.63 | +0.64 | +0.59 | +0.83 | +0.74 | +0.84 | +0.73 | +0.65 | +0.26 | $\pm 0.00$ |
| Y453 | +1.81 | +0.42 | +1.72 | +1.54 | +0.31 | +1.95 | +0.78 | +0.64 | +1.48 | +0.76 | +0.85 | +1.42 | +1.59 | +1.40 | +1.16 | +1.65 | +1.30 | +0.71 | +0.30 | $\pm 0.00$ |
| L455 | +1.62 | +0.33 | +2.09 | +1.96 | -0.14 | +1.89 | +0.61 | +0.13 | +1.69 | $\pm 0.00$ | +0.55 | +1.88 | +1.06 | +1.55 | +0.93 | +1.58 | +1.31 | +0.39 | -0.07 | -0.22 |
| F456 | +2.18 | +1.49 | +1.85 | +1.94 | $\pm 0.00$ | +2.45 | +1.20 | +1.09 | +1.61 | +1.25 | +1.30 | +1.83 | +2.51 | +1.68 | +1.17 | +2.12 | +1.66 | +1.32 | +0.52 | +0.57 |
| Y473 | +1.03 | +0.87 | +0.81 | +1.04 | +0.38 | +1.17 | +0.68 | +0.58 | +0.97 | +0.62 | +0.51 | +0.82 | +0.53 | +0.87 | +0.63 | +0.92 | +0.78 | +0.76 | +0.46 | $\pm 0.00$ |
| A475 | $\pm 0.00$ | +0.06 | +1.42 | +1.58 | -0.32 | +0.91 | +0.45 | +0.07 | +1.07 | +0.17 | +0.19 | +0.86 | +0.63 | +0.71 | +0.74 | +0.62 | +0.54 | +0.14 | -0.30 | -0.27 |
| G476 | +0.70 | +0.30 | +0.50 | +0.81 | +0.08 | $\pm 0.00$ | +0.19 | +0.44 | +0.63 | +0.47 | +0.29 | +0.37 | +1.03 | +0.59 | +0.52 | +0.50 | +0.44 | +0.40 | +0.17 | +0.23 |
| E484 | +0.10 | -0.04 | +0.07 | $\pm 0.00$ | -0.09 | +0.43 | +0.07 | -0.15 | +0.07 | +0.01 | +0.09 | +0.09 | -0.43 | +0.22 | +0.10 | +0.06 | -0.04 | -0.16 | -0.12 | -0.07 |
| F486 | +1.68 | +1.60 | +1.48 | +1.34 | $\pm 0.00$ | +1.89 | +0.84 | +0.90 | +1.08 | +0.74 | +0.85 | +1.33 | +0.81 | +1.20 | +0.95 | +1.46 | +1.23 | +0.93 | +0.31 | +0.50 |
| N487 | +1.13 | +0.19 | +0.97 | +1.19 | +0.02 | +0.75 | +0.18 | +0.51 | +1.08 | +0.39 | +0.25 | $\pm 0.00$ | +1.16 | +0.69 | +0.60 | +1.03 | +0.80 | +0.35 | +0.23 | -0.24 |
| Y489 | +2.82 | +2.59 | +2.79 | +2.20 | +1.38 | +3.43 | +1.74 | +1.93 | +1.98 | +2.01 | +1.68 | +2.58 | +2.84 | +1.97 | +1.85 | +2.87 | +2.47 | +2.36 | +0.84 | $\pm 0.00$ |
| Q493 | +1.52 | +0.83 | +1.46 | +1.36 | +0.10 | +1.84 | +0.42 | +0.56 | +0.97 | +0.45 | +0.57 | +1.15 | +1.36 | $\pm 0.00$ | +0.60 | +2.02 | +1.10 | +0.71 | -0.08 | +0.15 |
| G496 | +0.63 | +0.02 | +0.80 | +1.05 | +0.15 | $\pm 0.00$ | +0.33 | +0.41 | +1.12 | +0.58 | +0.40 | +0.67 | +1.11 | +0.78 | +0.71 | +0.44 | +0.41 | +0.22 | +0.18 | +0.21 |
| Q498 | +0.75 | -0.38 | +1.12 | +1.64 | -0.27 | +0.98 | +0.26 | -0.37 | +1.46 | -0.49 | -0.28 | +0.80 | +1.26 | $\pm 0.00$ | +0.84 | +0.74 | +0.19 | -0.01 | -0.25 | -0.26 |
| T500 | +1.62 | +1.57 | +1.08 | +1.93 | +0.36 | +2.30 | +0.91 | +0.80 | +1.74 | +0.40 | +0.46 | +1.01 | +1.75 | +1.47 | +0.99 | +1.38 | $\pm 0.00$ | +0.83 | +0.05 | +0.09 |
| N501 | +0.88 | +0.46 | +0.80 | +0.77 | -0.09 | +1.24 | +0.33 | +0.23 | +0.46 | +0.06 | -0.18 | $\pm 0.00$ | +1.19 | +0.56 | +0.43 | +1.25 | +0.42 | +0.20 | -0.08 | -0.08 |
| G502 | +1.21 | +0.91 | +1.38 | +2.16 | -0.51 | $\pm 0.00$ | +0.04 | +0.87 | +1.24 | +0.51 | +0.41 | +1.05 | +0.97 | +1.20 | +0.65 | +1.23 | +1.09 | +0.99 | -0.49 | -0.28 |
| V503 | +0.34 | +0.42 | +0.08 | +0.42 | +0.45 | +0.41 | +0.16 | +0.29 | +0.39 | +0.26 | +0.28 | +0.12 | +0.04 | +0.36 | +0.17 | +0.09 | +0.11 | $\pm 0.00$ | +0.14 | +0.44 |
| Y505 | +1.78 | +1.83 | +1.74 | +1.85 | +0.86 | +2.51 | +1.09 | +1.45 | +1.82 | +1.24 | +1.14 | +1.54 | +1.94 | +1.24 | +1.09 | +1.80 | +1.71 | +1.95 | +0.13 | $\pm 0.00$ |

a) no cumulative effects; free energy difference calculations for individual amino acid positions.
b) residues with a shortest atom distance below 5 Å between RBD and ACE2.

**Supplemental Table 2:** Effects of SARS-CoV2 RBD-exchanged amino acid residues on strengths of interaction with DPP-IV.

| amino acid<br>residues in<br>wt RBD <sup>b)</sup> | $\Delta\Delta G$ values in kcal / mol <sup>a)</sup> | | | | | | | | | | | | | | | | | | | |
| --- | --- | --- | --- | --- | --- | --- | --- | --- | --- | --- | --- | --- | --- | --- | --- | --- | --- | --- | --- | --- |
|  | A | C | D | E | F | G | H | I | K | L | M | N | P | Q | R | S | T | V | W | Y |
| R403 | +0.77 | +0.31 | +0.67 | +0.58 | +0.20 | +0.99 | +0.02 | +0.24 | +0.58 | +0.31 | +0.29 | +0.63 | +0.25 | +0.49 | $\pm 0.00$ | +0.64 | +0.38 | +0.53 | +0.11 | +0.31 |
| D405 | +0.43 | +0.45 | $\pm 0.00$ | +0.31 | -0.10 | +0.57 | -0.06 | +0.25 | +0.32 | +0.26 | +0.20 | +0.08 | +0.27 | +0.20 | +0.26 | +0.21 | +0.23 | +0.43 | -0.14 | +0.05 |
| E406 | +0.07 | +0.69 | +0.64 | $\pm 0.00$ | +0.02 | +0.78 | +0.08 | +0.50 | +0.72 | +0.27 | +0.16 | +0.43 | +1.13 | +0.47 | +0.38 | +0.67 | +0.61 | +0.39 | +0.19 | -0.01 |
| R408 | +0.53 | +0.35 | +0.11 | +0.19 | -0.11 | +0.51 | -0.13 | +0.46 | +0.16 | +0.07 | +0.03 | -0.04 | +0.11 | +0.18 | $\pm 0.00$ | +0.23 | +0.13 | +0.73 | +0.17 | +0.12 |
| Q409 | +1.05 | +0.64 | +0.64 | +0.62 | +0.06 | +0.66 | +0.11 | +0.23 | +0.47 | +0.39 | +0.38 | +0.48 | +0.81 | $\pm 0.00$ | +0.47 | +0.64 | +0.67 | +0.55 | +0.23 | +0.07 |
| T415 | +1.03 | +0.83 | +0.91 | +1.21 | +0.58 | +1.15 | +0.46 | +0.66 | +0.98 | +0.70 | +0.85 | +0.75 | +1.13 | +0.76 | +0.71 | +0.73 | $\pm 0.00$ | +0.60 | +0.59 | +0.69 |
| G416 | +0.75 | +0.34 | +0.74 | +1.49 | +0.44 | $\pm 0.00$ | +0.46 | +0.58 | +1.38 | +0.68 | +0.55 | +1.03 | +1.02 | +1.15 | +0.73 | +0.76 | +0.84 | +0.74 | +0.42 | +0.38 |
| K417 | +1.44 | +0.49 | +0.69 | +0.44 | -0.32 | +0.92 | +0.02 | +0.15 | $\pm 0.00$ | +0.07 | +0.10 | +0.64 | +0.45 | +0.50 | +0.46 | +1.23 | +0.91 | +0.90 | -0.34 | -0.18 |
| L455 | +1.17 | +0.56 | +1.30 | +1.16 | +0.37 | +1.54 | +0.64 | +0.40 | +0.99 | $\pm 0.00$ | +0.61 | +1.26 | +0.74 | +0.91 | +1.04 | +1.04 | +0.91 | +0.54 | +0.56 | +0.39 |
| F456 | +2.41 | +1.46 | +2.15 | +2.31 | $\pm 0.00$ | +2.56 | +1.42 | +1.09 | +1.90 | +1.09 | +1.39 | +1.84 | +2.50 | +1.85 | +1.65 | +2.19 | +1.74 | +1.24 | +0.60 | +0.75 |
| I472 | +1.07 | +0.64 | +1.28 | +1.30 | +0.29 | +1.12 | +0.69 | $\pm 0.00$ | +1.08 | +0.32 | +0.55 | +0.96 | +0.51 | +1.10 | +0.90 | +1.06 | +0.77 | +0.41 | +0.40 | +0.53 |
| Y473 | +1.33 | +1.00 | +1.31 | +1.48 | +0.41 | +1.50 | +0.88 | +0.55 | +1.22 | +0.63 | +0.53 | +1.17 | +0.90 | +1.15 | +0.87 | +1.24 | +0.94 | +0.63 | +0.51 | $\pm 0.00$ |
| Q474 | +0.14 | -0.01 | +0.20 | +0.38 | -0.58 | +0.11 | -0.26 | -0.35 | +0.27 | -0.53 | -0.12 | +0.16 | -0.27 | $\pm 0.00$ | -0.18 | +0.19 | +0.18 | -0.27 | -0.37 | -0.36 |
| A475 | $\pm 0.00$ | -0.12 | +1.93 | +1.89 | -1.14 | +1.15 | +0.25 | -0.05 | +1.07 | -0.09 | -0.23 | +1.35 | +1.20 | +1.10 | +0.56 | +1.05 | +0.91 | +0.30 | -1.26 | -1.14 |
| G476 | +0.58 | -0.88 | +0.75 | +1.40 | -1.61 | $\pm 0.00$ | -0.81 | -0.69 | +1.16 | -0.41 | -0.59 | +0.40 | +1.11 | +0.63 | +0.14 | +0.22 | +0.23 | -0.51 | -0.99 | -0.98 |
| S477 | +0.14 | -0.03 | +0.36 | +0.59 | +0.32 | +0.33 | +0.28 | +0.20 | +0.56 | +0.21 | +0.21 | +0.39 | -0.11 | +0.42 | +0.47 | $\pm 0.00$ | +0.13 | +0.15 | +0.36 | +0.36 |
| T478 | +0.31 | +0.28 | +0.55 | +1.05 | +0.15 | +0.60 | +0.16 | +0.02 | +1.00 | -0.02 | +0.08 | +0.44 | +0.36 | +0.65 | +0.69 | +0.17 | $\pm 0.00$ | -0.14 | +0.51 | +0.38 |
| P479 | +0.25 | +0.35 | +0.19 | +0.41 | +0.51 | +0.50 | +0.30 | +0.26 | +0.41 | +0.29 | +0.38 | +0.26 | $\pm 0.00$ | +0.37 | +0.36 | +0.14 | +0.14 | +0.25 | +0.50 | +0.38 |
| C480 | +0.38 | $\pm 0.00$ | +0.60 | +0.67 | +0.14 | +0.89 | +0.43 | +0.16 | +0.49 | +0.17 | +0.26 | +0.71 | +0.46 | +0.53 | +0.47 | +0.36 | +0.43 | +0.33 | +0.13 | +0.08 |
| V483 | +0.41 | +0.16 | +0.53 | +0.24 | +0.31 | +0.54 | +0.23 | +0.05 | +0.14 | +0.20 | +0.17 | +0.36 | +0.87 | +0.11 | +0.19 | +0.37 | +0.15 | $\pm 0.00$ | +0.44 | +0.35 |
| E484 | -0.20 | -0.21 | 0.06 | $\pm 0.00$ | -0.32 | +0.36 | -0.08 | -0.34 | -0.05 | -0.40 | -0.22 | -0.02 | -0.22 | -0.03 | -0.11 | 0.00 | -0.01 | -0.25 | -0.27 | -0.22 |
| G485 | +1.24 | +0.45 | +0.83 | +1.28 | -0.00 | $\pm 0.00$ | +0.45 | +0.45 | +0.80 | +0.33 | +0.10 | +0.19 | +0.53 | +0.98 | +0.52 | +0.86 | +0.60 | +0.41 | +0.24 | +0.50 |
| F486 | +4.18 | +3.34 | +3.45 | +3.20 | $\pm 0.00$ | +4.41 | +2.05 | +2.51 | +3.48 | +2.21 | +2.09 | +3.26 | +3.46 | +2.88 | +2.50 | +3.90 | +3.24 | +3.13 | +0.87 | +0.82 |
| N487 | +0.42 | -1.78 | +1.39 | +2.62 | -2.27 | -0.12 | -0.83 | -1.11 | +2.05 | -1.06 | -1.38 | $\pm 0.00$ | +0.88 | +0.74 | +0.10 | +0.39 | +0.40 | -0.89 | -2.03 | -2.25 |
| C488 | +2.03 | $\pm 0.00$ | +2.49 | +2.74 | +0.30 | +2.23 | +1.19 | +1.06 | +2.25 | +0.99 | +0.83 | +2.08 | +3.53 | +1.55 | +1.53 | +1.90 | +2.02 | +1.33 | +0.46 | +0.40 |
| Y489 | +2.37 | +2.04 | +2.53 | +1.97 | +0.81 | +2.76 | +1.32 | +1.49 | +1.74 | 1.75 | +1.33 | +2.15 | +2.53 | +1.71 | +1.41 | +2.45 | +2.16 | +1.90 | +0.58 | $\pm 0.00$ |
| Q493 | +0.71 | +52 | 0.84 | +0.80 | -0.13 | +0.98 | +0.27 | +0.15 | +0.83 | +0.04 | +0.31 | +0.58 | +0.87 | $\pm 0.00$ | +0.63 | +1.02 | +0.47 | +0.22 | -0.07 | +0.11 |
| T500 | +0.17 | -0.02 | -0.15 | +0.12 | +0.19 | +0.26 | +0.10 | +0.26 | +0.12 | +0.14 | +0.17 | -0.09 | +0.33 | +0.20 | +0.18 | +0.03 | $\pm 0.00$ | +0.26 | +0.15 | +0.15 |
| N501 | +0.67 | +0.73 | +0.67 | +0.38 | +0.30 | +0.84 | +0.02 | +0.44 | +0.21 | +0.43 | +0.35 | $\pm 0.00$ | +0.68 | +0.21 | +0.18 | +0.79 | +0.09 | +0.46 | +0.25 | +0.19 |
| G502 | +1.59 | +0.92 | +1.91 | +2.50 | +0.19 | $\pm 0.00$ | +0.55 | +1.27 | +2.13 | +0.88 | +0.91 | +1.68 | +1.15 | +1.89 | +1.44 | +1.50 | +1.38 | +1.37 | +0.07 | +0.46 |
| V503 | +0.27 | +0.46 | +0.01 | +0.37 | +0.48 | +0.42 | +0.10 | +0.23 | +0.25 | +0.17 | +0.18 | +0.05 | -0.17 | +0.24 | +0.02 | +0.10 | +0.05 | $\pm 0.00$ | +0.25 | +0.36 |
| Y505 | +1.85 | +2.07 | +1.71 | +1.67 | +0.71 | +2.35 | +1.20 | +1.45 | +1.72 | +1.11 | +0.94 | +1.41 | +1.94 | +1.20 | +1.32 | +1.70 | +1.64 | +1.57 | +0.48 | $\pm 0.00$ |

a) no cumulative effects; free energy difference calculations for individual amino acid positions.
b) residues with a shortest atom distance below 5 Å between RBD and DPP-IV.

## Notes

1) The authors declare no conflicts of interest. (e.g., pharmaceutical stock ownership, consultancy, advisory board membership, relevant patents, or research funding)

### Competing Interest Statement

The authors have declared no competing interest.

